# Septal HDAC1 facilitates long-lasting extinction of social fear in male mice

**DOI:** 10.1101/2020.09.05.284406

**Authors:** Anna Bludau, Andreas Paul, Inga D Neumann, Rohit Menon

**Affiliations:** Department of Behavioural and Molecular Neurobiology, Regensburg Center of Neuroscience; University of Regensburg, Regensburg, Germany

## Abstract

Social anxiety disorder (SAD) is primarily caused by traumatic social events and is characterized by intense fear and avoidance of socially enriched contexts. Formation of such intense social aversion requires learning of the association between trauma and originally neutral social stimuli. Acetylation of histones in response to learning is critical for long-term memory formation. Class I histone deacetylases (HDACs) have emerged as powerful negative regulators of long-term memory formation. However, the lack of clarity in isoform and spatial specificity in HDAC function exists. Utilizing the social fear conditioning (SFC) as well as cued fear conditioning (CFC) paradigms, we aimed to functionally characterize the role of class I HDACs, especially HDAC1, in the regulation of aversive memories in social *versus* non-social contexts. Using cFos expression as a marker of cellular activity, we revealed that the lateral septum (LS) was specifically activated post-acquisition in socially fear conditioned (SFC^+^) mice. We further measured an increase in activity-inducing HDAC1 phosphorylation at serine residues within the septum of SFC^+^ mice during extinction of social fear. Pharmacological inhibition of HDAC1 within the LS facilitated, while viral overexpression of septal HDAC1 impaired extinction of social fear. Finally, we found that the abovementioned facilitation of extinction memory by HDAC1 inhibition was long-lasting and persists even after 30 days of extinction. Our results show that LS-HDAC1 is a key regulator of long-lasting social fear extinction, which points towards HDAC1 as a potential therapeutic target for SAD.

## Introduction

Fear is an essential and highly adaptive behavioral response. However, its post-stimulus persistence is characteristic of anxiety disorders. Social anxiety disorder (SAD), which is the second most prevalent anxiety disorder, is characterized by intense fear and avoidance of social situations [1,2]. One of the most prominent therapeutic interventions for SAD consists of exposure-based cognitive behavioural therapy in combination with pharmacotherapy with benzodiazepines or selective serotonin reuptake inhibitors. However, SAD patients are often treatment-resistant or report a re-emergence of previously extinguished fear responses [2–4], which might partially originate from insufficient consolidation of extinction memory [5]. Although there has been a significant increase in our understanding of the molecular processes underlying extinction memory consolidation in non-social contexts [6], mechanisms regulating social fear extinction are rather unknown with the consequence that efficient and specific therapeutic options for SAD are missing. So far, a major hurdle in developing specific treatment options for SAD has been the lack of animal models resembling its symptoms, i.e., the intense fear and avoidance of conspecifics. To this end, we established the social fear conditioning (SFC) paradigm in mice, which is based on operant fear conditioning principles and generates robust social fear [7]. Using this model, we have previously identified the lateral septum (LS) - a subcortical limbic region – as being dynamically involved in the extinction of SFC-induced fear of conspecifics in both male [8] and female [9] mice. Using the SFC paradigm we endeavour to further our understanding of epigenetic mechanisms regulating social fear extinction.

Concurring evidence suggests the importance of post-translational acetylation of N-terminal tails of histone residues in consolidation and maintenance of fear memory [10]. Histone deacetylases (HDACs), which catalyse the removal of acetyl groups from these N-terminal tails of histones, are key regulators of gene expression and have been found to alter social behaviour [11] as well as learning and memory [12]. Several isoforms of HDACs are found in rodents and humans, and they show remarkable brain region and task-dependent functional specificity [13]. For instance, hippocampal HDAC1 was found to enhance associative memory in a contextual fear conditioning paradigm [14]. Hippocampal HDAC2, on the other hand, has been functionally linked to reduced synaptic plasticity and impairment in long-term object and spatial memory formation [15,16]. Similar studies have implicated differential contribution of HDAC3 in the formation of robust aversive contextual and cued fear memories in the hippocampus and basal amygdala, respectively [17]. Taking both, their role in regulating fear learning as well as their sensitivity to pharmacological manipulation, into account, factors regulating histone acetylation present a novel avenue for treating anxiety disorders. Hence in recent years, many studies have used HDAC inhibitors as cognitive enhancers to rescue learning deficits [18–20]. However, the contribution of HDACs in fear memory regulation has, so far, exclusively been studied in non-social models of fear conditioning, while their contribution in the regulation of social fear remains elusive.

Here, we studied the contribution of epigenetic mechanisms, specifically of HDAC1-mediated regulation of fear extinction within the LS towards social (SFC) and non-social (cued fear conditioning; CFC) fear in male mice. Using a combination of molecular, pharmacological and viral approaches we show that HDAC1 within the LS negatively regulates long-lasting fear extinction memory in a social context.

## Materials and Methods

### Animals

Male CD1 mice (Charles River Germany, 8-11weeks of age at the start of experiments) were kept group-housed under standard laboratory conditions (12/12h light/dark cycle, lights on at 06:00, 22°C, 60% humidity, food and water *ad libitum*) in polycarbonate cages (16 x 22 x 14 cm) until 3 days before each behavioural experiment. Age and weight-matched male CD1 mice were used as social stimuli in the SFC paradigm. All experimental procedures were performed between 08:00 and 12:00hrs in accordance with the Guide for the Care and Use of Laboratory Animals of the Government of Oberpfalz, ARRIVE guidelines [21] and the guidelines of the NIH.

### Experimental protocols

#### Experiment 1: Effect of SFC and CFC on neuronal activity (Figure 1)

To compare neuronal activation patterns in response to SFC and CFC, mice were transcardially perfused either on day 1 of the SFC paradigm with (SFC^+^/Ext^-^) or without (SFC^-^/Ext^-^) social fear acquisition, or on day 2 after social fear extinction training (SFC^+^/Ext^+^ and SFC^-^/Ext^+^), or at identical time points in the CFC paradigm (day1; post-acquisition: CFC^+^/Ext^-^ and CFC^-^/Ext^-^; day 2; post-extinction: CFC^+^/Ext^+^ and CFC^-^/Ext^+^). Mice were killed 90min after respective behavioural procedures. Perfused brains were processed, immunohistochemically stained for cFos, and the number of immuno-positive cells per 0.5mm^2^ was counted in the dorsal and ventral LS as well as the basolateral amygdala (BLA) and paraventricular nucleus of the hypothalamus (PVN) [22].

**Figure 1.**
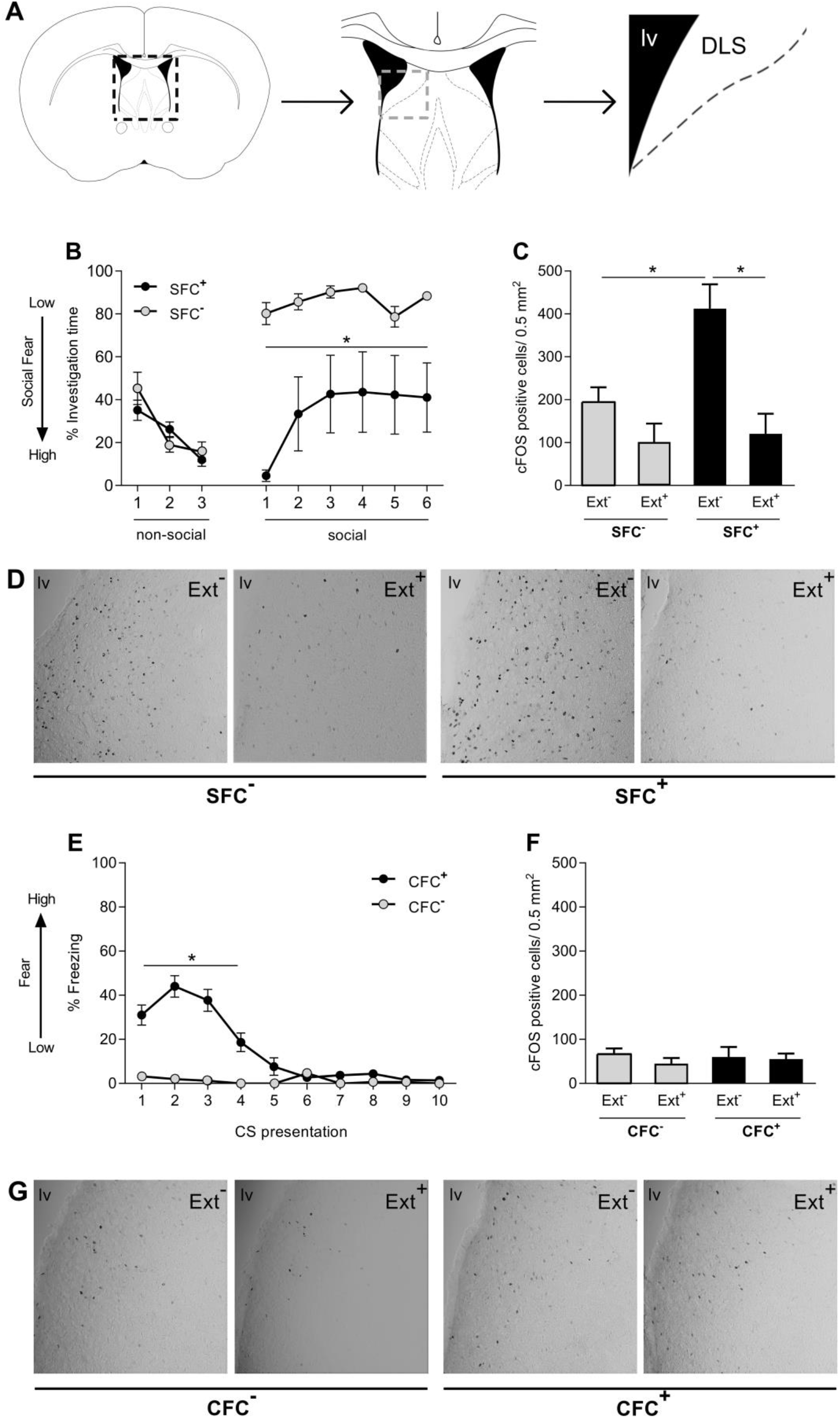
Social fear conditioning (SFC) leads to activation of the dorsal lateral septum (LS). Numbers of cFos-immunopositive cells were counted after social fear conditioning (SFC; B-D) and cued fear conditioning (CFC; E-G) within the dorsal LS (Bregma 0.86). Mice were sacrificed 90 min after either social fear acquisition (SFC^-^/Ext^-^ and SFC^+^/Ext^-^; n = 6), cued fear acquisition (CFC^-^/Ext^-^ and CFC^+^/Ext^-^; n = 6), social fear extinction (A; SFC^-^/Ext^+^ and SFC^+^/Ext^+^; n = 6) or cued fear extinction (E; CFC^-^/Ext^+^ and CFC^+^/Ext^+^; n = 6). Data represent mean percentage of investigation time ± SEM (B), mean number of cFos-immunopositive neurons/ 0.5 mm^2^ + SEM (C-D and F-G), and mean percentage of CS-elicited freezing ± SEM (E). *p<0.05 *vs* SFC^-^ (B), indicated groups (C) or CFC^-^ (E).

#### Experiment 2: Effect of social fear extinction on HDAC1 protein levels and phosphorylation within the septum (Figure 2)

To test for the influence of extinction on HDAC1 protein level and phosphorylation within the septum, mice were rapidly decapitated under anaesthesia on day 2 of the SFC paradigm at different time points of the social fear extinction protocol, i.e. either at 90 min directly after exposure to the 1^st^ social stimulus in an abridged social fear extinction protocol, or 90 min after exposure to the 6^th^ social stimulus, i.e. after termination of the standard procedure of social fear extinction. Septal tissue micro punches were obtained, and local HDAC1 protein levels and phosphorylation was measured in nuclear extracts using Western blot.

**Figure 2.**
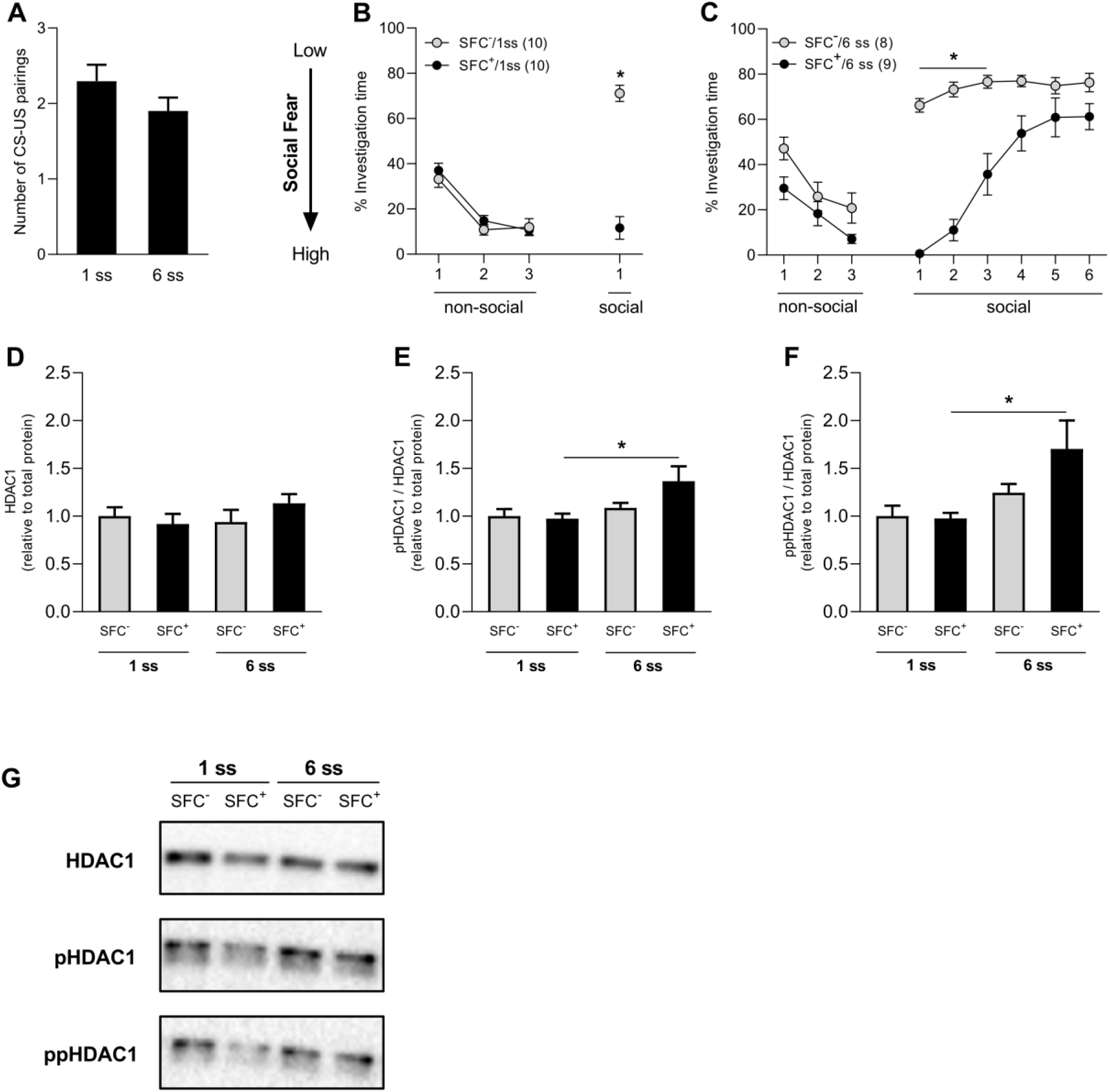
HDAC1 phosphorylation on serine residues is upregulated in response to social fear extinction within the septum. Mice were social fear conditioned on day 1 of the SFC paradigm. Number of CS-US pairings for both groups of SFC^+^ mice, i.e., those exposed to 1 social stimulus (1 ss) and 6 social stimuli (6 ss) during extinction (A). Total protein was isolated from mice septum 90 min either after abridged social fear extinction (SFC^-^/1 ss and SFC^+^/6 ss n = 10/group; B) or complete extinction (SFC^-^/6 ss and SFC^+^/6 ss; n = 8-9/group; C). Relative protein levels of HDAC1 (D), pHDAC1 (Ser421) (E) and ppHDAC (Ser421, Ser423) (F) were measured by Western blot. Representative western blot images for HDAC1, pHDAC1 and ppHDAC1 (G). Data represent mean CS-US pairings + SEM (A), mean investigation time ± SEM (B and C), or fold change + SEM (D-F). *p<0.05 SFC^+^ *vs* SFC^-^(B and C) or between indicated groups (E and F).

#### Experiment 3: Effect of pharmacological blockade of HDAC1 on extinction of social fear (Figure 3)

To assess the causal role of HDAC1 blockade in social fear extinction, MS275 (an *ortho*-amino anilide based small molecule inhibitor with high affinity for HDAC1) [23] or vehicle (Veh) was infused bilaterally into the LS 120min prior to the start of social fear extinction training leading to the following groups: SFC^+^/Veh, SFC^+^/MS275, SFC^-^/Veh, and SFC/MS275. The time mice spent investigating the social stimuli was recorded as indicator of social fear.

**Figure 3.**
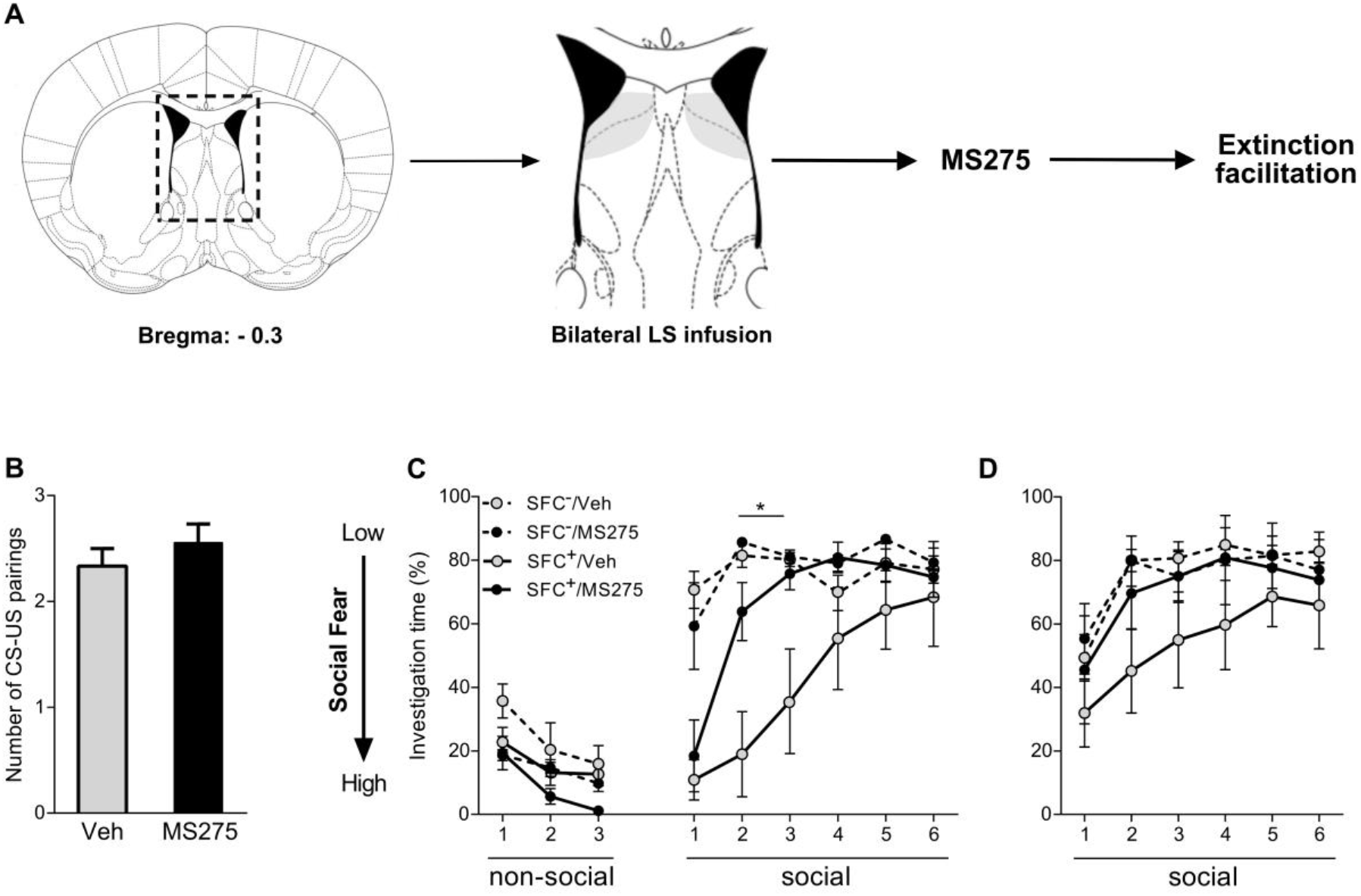
Pharmacological blockade of HDAC1 within the lateral septum (LS) specifically facilitates extinction of social fear in male mice. Schematic representation of LS including pharmacological interventions. Infusions within the highlighted area were considered as correct cannula placements (A). Number of CS-US pairings conditioned (SFC^+^) mice during acquisition of social fear on day 1 (B). Percentage of investigation time of three non-social (empty cages) and six social (cage with a conspecific) stimuli during social fear extinction (Day 2; C) and social fear extinction recall (Day 3; D) of unconditioned (SFC^-^) and SFC^+^ mice. SFC^+^ and SFC^-^ mice were bilaterally infused with vehicle (Veh; Ringer; 0.2 μl/hemisphere) or MS275 (375 ng/0.2 μl/hemisphere) into the LS 120 min prior to extinction training. Data represent mean CS-US pairings + SEM (B) or mean percentage investigation time ± SEM (C-D). *p<0.05 for SFC^+^/Veh *vs* SFC^+^/MS275, SFC^-^/Veh, and SFC^-^/MS275.

#### Experiment 4: Effect of viral overexpression of septal HDAC1 on extinction of social fear (Figure 4)

To assess the effect of septal HDAC1 overexpression on extinction of social fear, adenoviral vector (AAV)-mediated overexpression of HDAC1 or a GFP reporter (i.e., control vector) was performed by infusing respective vectors into the septum of mice 3 weeks prior to the start of the SFC experiment leading to the following groups: SFC^+^/GFP, SFC^+^/HDAC1, SFC^-^/GFP, and SFC^-^/HDAC1. The time mice spent investigating the social stimuli during social fear extinction was recorded as indicator of social fear.

**Figure 4.**
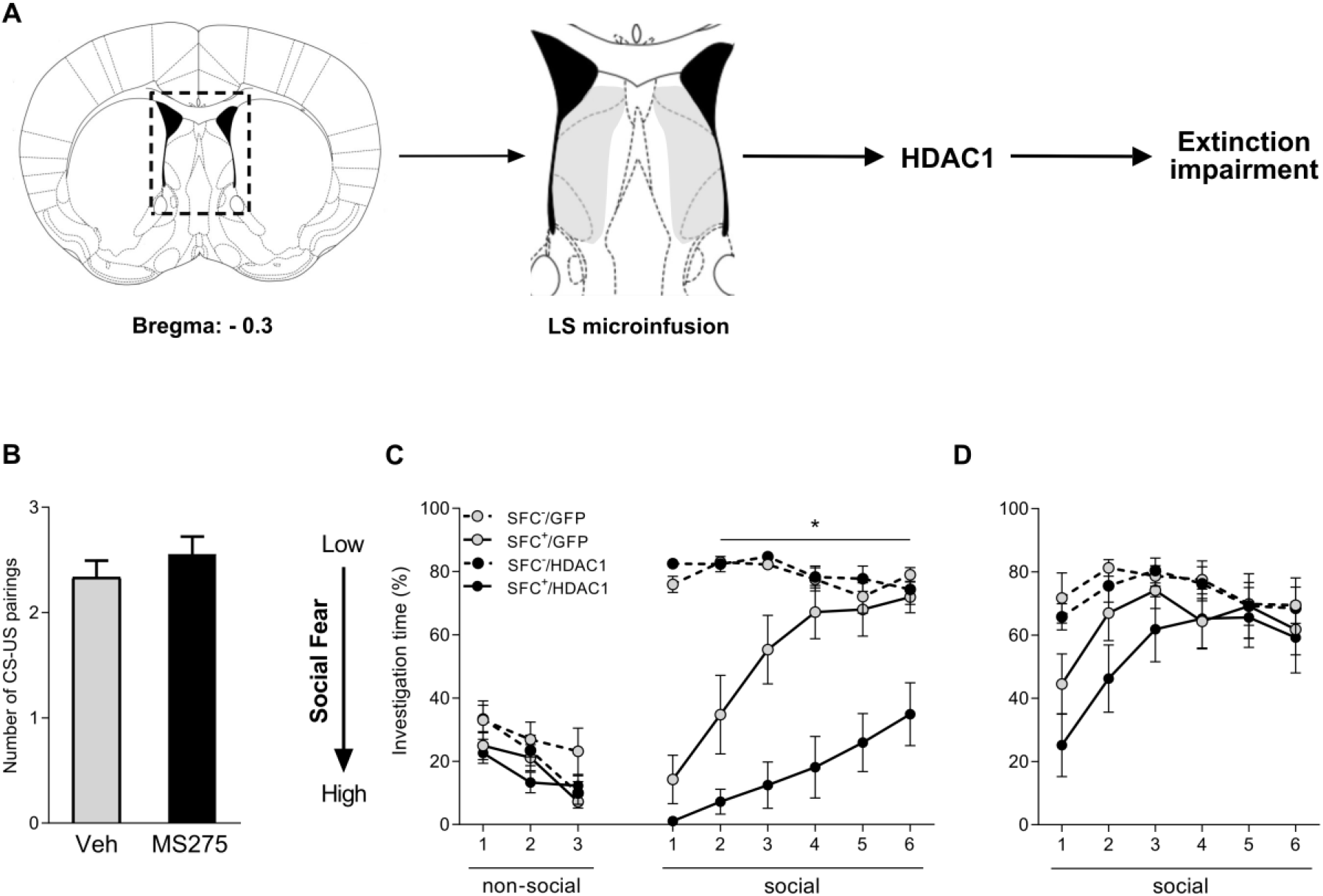
Viral overexpression of HDAC1 within the lateral septum impairs social fear extinction in male mice. Schematic representation of the septum including viral manipulations (A). The highlighted area was considered as adequate infusion localization. Male mice were injected with AAV-hSyn-HDAC-GFP-WPRE (HDAC1) or AAV-hSyn-GFP-WPRE (GFP; control) 3 weeks prior to acquisition of social fear. Number of CS-US pairings of conditioned (SFC^+^) mice during acquisition of social fear on day 1 (B). Percentage of investigation time of three non-social (empty cages) and six social (cage with a conspecific) stimuli during social fear extinction (Day 2; C) and social fear extinction recall (Day 3; D) in unconditioned (SFC^-^) and SFC^+^ mice. Data represent mean CS-US pairings + SEM (B) or mean percentage investigation time ± SEM (C-D). +p<0.05 SFC^+^/HDAC1 *vs* SFC^+^/GFP (C).

#### Experiment 5: Effect of pharmacological blockade of HDAC1 on consolidation of social fear extinction memory (Figure 5)

To assess the effect of HDAC1 blockade on long-term consolidation of social fear extinction memory, SFC^+^ mice were infused with MS275 or Veh bilaterally in the LS of conditioned mice 120min prior to social fear extinction training on day 2 of the SFC paradigm, resulting in the following groups: SFC^+^/Veh and SFC^+^/MS275. To test for long-term consolidation of extinction memory, recall was performed 30 days after extinction training. The time mice spent investigating the social stimuli was recorded as indicator of social fear.

**Figure 5.**
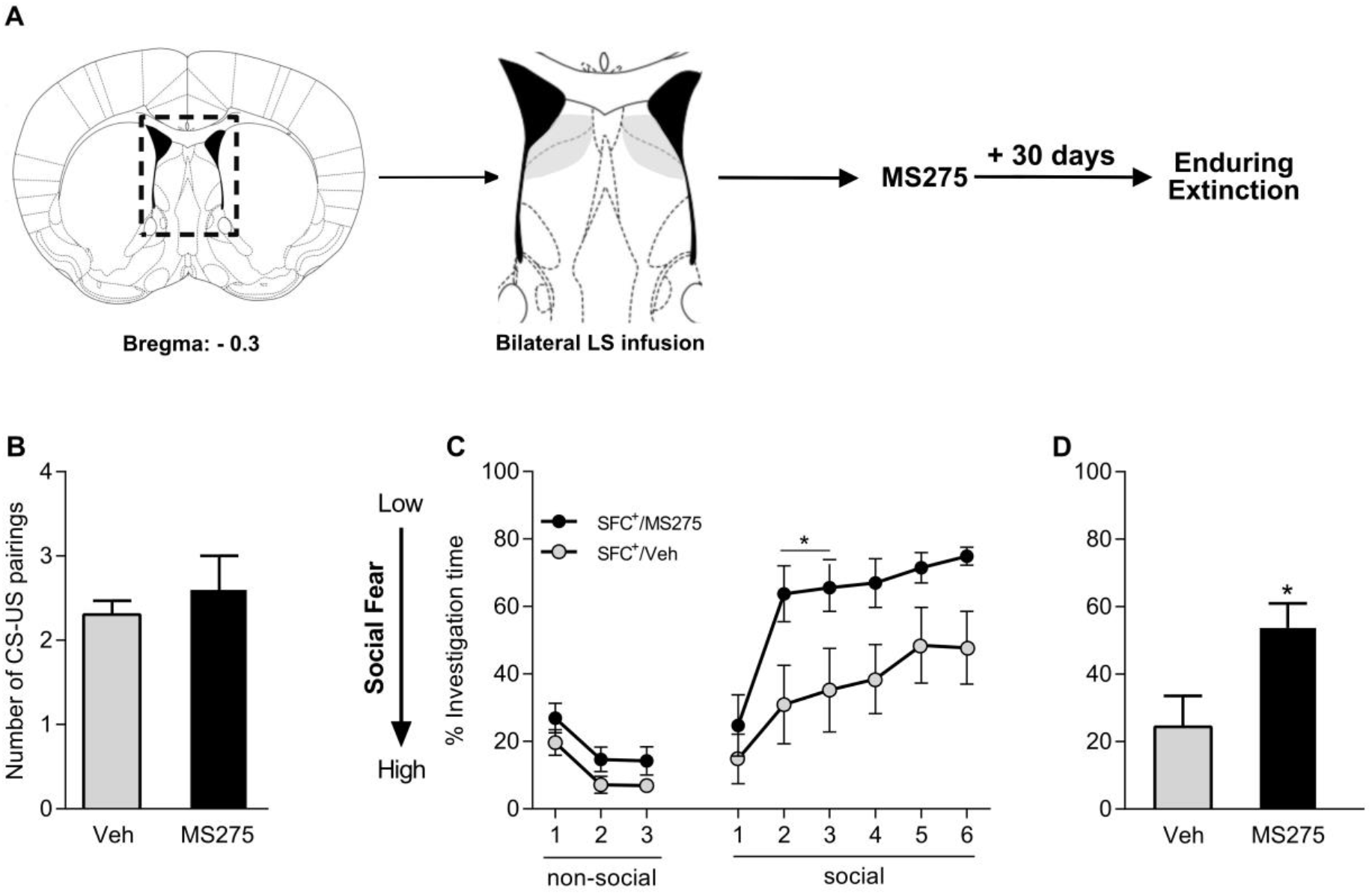
Pharmacological blockade of HDAC1 within the lateral septum (LS) leads to enhanced long-lasting consolidation of social fear extinction memory in male mice. Schematic representation of LS including pharmacological interventions (A). Infusions within the highlighted area were considered as correct cannula placements. CS-US pairings of conditioned (SFC^+^) mice during acquisition of social fear on day 1 (B). Percentage of investigation time of three non-social (empty cages) and six social (cage with a conspecific) stimuli during social fear extinction (day 2; C) and social fear extinction recall (day 31; D) of SFC^+^ mice, which were bilaterally infused with vehicle (Veh; Ringer; 0.2μl/hemisphere) or MS275 (375ng/0.2μl/hemisphere) into the LS 120min prior to extinction training. Data represent mean CS-US pairings + SEM (B) or mean percentage investigation ± SEM (C-D). *p<0.05 SFC^+^/MS275 *vs* SFC^+^/Veh (C and D).

### SFC paradigm

SFC was performed as previously described with minor modifications for analysis of long-lasting extinction [7]. For details see supplementary methods.

### CFC paradigm

CFC was performed as described previously [7]. For details see supplementary methods.

### Stereotaxic implantations

Guide cannulas (23G, 8mm length; Injecta GmbH) for bilateral drug infusion into the LS were implanted stereotaxically with the tip resting 2mm above the LS target region (from bregma: anteriorposterior - 0.3mm, mediolateral ±0.5mm, dorsoventral −1.6mm; Paxinos and Franklin, 2001) under isoflurane anesthesia (Forene, Abbott GmbH) [8,22]. To avoid postsurgical infections and pain, all mice received a subcutaneous injection of antibiotics (10mg/kg Baytril, Bayer GmbH) and analgesics (0.05mg/kg Buprenorphine, Bayer). In order to reduce post-surgical stress, all mice were handled once a day for at least five days before experiments.

### Intra-septal drug infusions

Mice received bilateral infusions of either MS275 (375ng/0.2μl/hemisphere) or Veh (0.2μl/hemisphere; 0.5% DMSO in sterile Ringer solution) into the LS via a 27G infusion cannula inserted into the implanted guide cannulas connected to a 10-μl Hamilton syringe. All infusions were performed 120min before SFC or CFC extinction training. The dose and timing of MS275 were selected based on a previous study [14]. The correct infusion sites were histologically verified, and animals with misplaced infusions were excluded from statistical analysis.

### Intra-septal viral infusions

The AAV (Vector Biolabs) for overexpression of HDAC1, i.e., AAV-hSyn-HDAC1-GFP-WPRE (HDAC1; 2.5 x 10^13^ GC/ml;), or the control vector, i.e., AAV-hSyn-GFP-WPRE (GFP; 2.5 x 10^13^ GC/ml;), were bilaterally infused into the septum (anterior-posterior from bregma +0.3mm, lateral ±0.5mm; dorsoventral - 3.8mm, −3.4mm, −3.1mm, −2.8 mm) with four 70-nl infusions/hemisphere. Infusions were performed 3 weeks prior to the start of behavioral experiments as previously described [9].

### cFos immunohistochemistry

Mice were transcardially perfused with 4% paraformaldehyde under deep anesthesia. Brains were removed and further processed to analyze cFos immunoreactivity as previously described [22]. Briefly, brain slices were incubated with a polyclonal primary antibody (cFos: 1:4000; sc-52, Santa Cruz) for 24hrs and a biotinylated goat-anti-rabbit secondary antibody (1:1500; Vector Laboratories) for 90min. Thereafter, cFos-positive neurons were visualised using an avidin-biotin-horseradish peroxidase procedure (ABC-Kit, Vector Laboratories) with 3, 3’-diaminobenzidine as the chromogen. Cells that were stained with a nuclear brown-black reaction product were considered as cFos-positive. A stereotaxic atlas (Paxinos and Franklin, 2^nd^ edition, 2001) was used to anatomically localize labelled cells. The number of cFos-positive cells was counted bilaterally in 3 to 4 sections depicting the dorsal and ventral LS (Bregma: 0.1mm to −0.10mm), BLA (Bregma: −0.8mm to −1.8mm), and PVN (Bregma: - 0.5mm to −1.2mm) in a tissue area of 0.5mm^2^ by an observer blind to the experimental groups. The average of these cell counts was calculated for each animal.

### Protein extraction and analysis

Mice were subjected to SFC as mentioned above and left undisturbed in their home cages for 90min after behavioural training. Mice were then sacrificed under anaesthesia, their brains were rapidly removed, and the septum (Bregma: 0.1mm to −0.10mm) was dissected. Subsequently, nuclear proteins were extracted using the EpiQuick Nuclear Extraction Kit (Epigentek) according to the manufacturer’s protocol.

Western blot analysis was used to detect HDAC1 expression and phosphorylation levels. 15μg of protein were resolved onto 12% TGX Stain-Free Gels (Bio-Rad) and transferred to nitrocellulose membranes. In all cases, the membranes were blocked with 5% milk, 0.05% Tween-20 in Tris-buffered saline followed by overnight incubation with the following antibodies: rabbit-anti-HDAC1 (Thermo Fisher; PA1-860; 1:2000), HDAC1 phosphorylation on serine residue 421 (rabbit-anti-pHDAC1(S421); Thermo Fisher; PA5-36810; 1:2000), as well as phosphorylation on serine 421 and 423 (rabbit-anti-pHDAC1(S421, S423), Thermo Fisher; PA5-36911; 1:2000). Immunolabeled bands were detected using the HRP-conjugated goat-anti-rabbit antibody (Cell Signaling; #7074; 1:1000) following an enhanced chemiluminescence system (SuperSignal West Dura, Thermo Fisher). Total protein was used as internal loading control.

### Scoring of behavior and statistical analysis

Social investigation times as indicator of social fear were manually scored by an observer blind to treatment using JWatcher (1.0, Macquarie University and UCLA). In the CFC paradigm, the CS-elicited freezing was considered as cued fear expression and measured using computerized scoring (TSE Systems GmbH). For statistical analysis, GraphPad Prism version 6.0 for Windows (GraphPad Software) was used. Detailed report for all statistical analysis is available in table 1 (supplementary information). All figures were produced using Affinity designer.

## Results

### Experiment 1: The LS is specifically activated by SFC, but not CFC

Successful social fear conditioning of SFC^+^ mice was indicated by reduced investigation of the social stimuli during social fear extinction training on day 2 (p<0.05 vs SFC^-^; Figure 1B). Similarly, successful cued fear conditioning of CFC^+^ mice was indicated by increased freezing in response to the CS-presentations on day 2 (p<0.05 vs. CFC^-^ mice; Figure 1E).

A significant increase in the number of cFos-positive cells was seen within the dorsal LS of SFC^+^/Ext^-^ mice 90 min after social fear acquisition (p<0.05 vs SFC^-^/Ext^-^), which returned to baseline levels 90 min after extinction training (p<0.05 vs SFC^+^/Ext^+^) (Figure 1C). A comparable dynamic neuronal response within the dorsal LS was not observed in response to cued fear acquisition or extinction (Figure 1F). In contrast to the dorsal LS, the ventral LS did not show differential activation in response to social fear acquisition, as there was no post-acquisition difference in the number of cFos-expressing cells within the ventral LS of SFC^-^ and SFC^+^ mice (Supplementary Figure S2A). However, the number of cFos-positive cells was significantly reduced post-extinction in both SFC^-^ and SFC^+^ mice in comparison to their respective post-acquisition groups (Supplementary Figure S2A). As the BLA is an established neuronal correlate of fear expression in mice [24,25], neuronal activation within the BLA in response to both SFC and CFC was measured as a positive control. Thus, an increased number of cFos-immunopositive cells in the BLA of conditioned mice was observed 90 min after both social (p<0.05 vs SFC^-^/Ext^-^; Supplementary Figure S2B) and cued (p<0.05 vs CFC^-^/Ext^-^; Supplementary Figure S2E) fear acquisition. Activity within the PVN was analyzed as a positive control for stress response in both SFC and CFC. PVN exhibited post-acquisition enhancement in activation in both SFC^+^ (p<0.05 vs SFC^-^/Ext^-^; Supplementary Figure S2C) and CFC^+^ (p<0.05 vs CFC^-^/Ext^-^; Supplementary Figure 2F) mice in comparison to their unconditioned counterparts suggesting that acquisition of fear, irrespective of the nature of the US is a stressful process.

### Experiment 2: HDAC1 phosphorylation on serine residues is upregulated in response to social fear extinction in the septum of fear conditioned mice

Class I HDACs are heavily implicated in regulation of memory formation and HDAC1 is especially implicated in regulation of contextual fear memories [14]. Amongst all class I HDACs (i.e. HDAC1, HDAC2, HDAC3 and HDAC8) we found *Hdac1* mRNA levels to be dynamically regulated within the septum in response to SFC, with its expression being the highest at 90 min after acquisition in SFC^+^ mice (Supplementary Figure S2B).

Therefore, we analyzed HDAC1 protein levels of septal tissue micro punches (*above different*) after exposure to either the 1^st^ social stimulus (1 ss) or 6^th^ (6 ss) social stimuli during social fear extinction as explained in the experimental protocols. Firstly, the number of CS-US pairings (Figure 2A) was similar between the 2 social exposure SFC^+^ groups. However, SFC^+^ mice spent significantly less time investigating the single (Figure 2B; p<0.05) or repeated social stimuli (Figure 2C; p<0.05) compared to SFC^-^ animals, depicting higher levels of social fear. Western blot analysis did not reveal differences in HDAC1 protein levels within the septum at 1 ss or 6 ss during extinction of social fear (Figure 2D). As post-translational phosphorylation of HDAC1 on ser421, i.e., pHDAC1, and double-phosphorylation at ser421 and ser423, i.e., ppHDAC1, are known to increase the deacetylase activity of HDAC1 [26], we quantified pHDAC1 and ppHDAC1 levels within the septum to assess the activation of HDAC1 as factor of social fear extinction. Both pHDAC1 and ppHDAC1 levels were increased in SFC^+^ mice, which underwent the standard extinction training and were exposed to 6 social stimuli (6ss), in comparison to those, which were only exposed to 1 ss during extinction (Figure 2E; p<0.05 for pHDAC1 and Figure 2F; p<0.05 for ppHDAC1), suggesting an increase in HDAC1 activity throughout extinction of social fear. Moreover, ppHDAC1 levels negatively correlated with the fold-change (calculated as the ratio of the average social investigation during exposure to social stimuli 5 and 6 to the average of social investigation during exposure to social stimuli 1 and 2) in social investigation throughout extinction of social fear in both SFC^-^ (Supplementary Figure S2H; p<0.05) and SFC^+^ (Supplementary Figure S2K; p<0.05) mice.

### Experiment 3: Pharmacological inhibition of LS-HDAC1 facilitates extinction of social fear

Based on the elevated HDAC1 phosphorylation within the septum after extinction of social fear in SFC^+^ mice, we studied the behavioral consequences of local inhibition of HDAC1 120 min prior to social fear extinction. For comparison, HDAC1 inhibition was also performed prior to cued fear extinction (see Supplementary Figure S3). During acquisition of social fear (Figure 3B), all mice received the same number of CS-US pairings. On day 2, all SFC^+^ mice infused with either MS275 or Veh in the LS expressed high levels of social fear during the presentation of the first social stimulus, as reflected by reduced social investigation. However, social fear extinction was accelerated by MS275-treatment, as the SFC^+^/MS275 group already displayed an elevated social investigation time during exposure to the 2^nd^ and 3^rd^ social stimulus (p<0.05 vs SFC^+^/Veh; Figure 3C). In SFC^-^ mice, treatment with MS275 did not affect social investigation times (Figure 3C–3D). All groups showed similarly high levels of social investigation during social fear recall on day 3 of the SFC paradigm (Figure 3D) indicating successful extinction of social fear. In contrast to SFC, MS275-treatment prior to cued fear extinction did not affect the freezing behavior of CFC mice (Supplementary Figure S3C).

### Experiment 4: AAV-mediated overexpression of HDAC1 within the LS impairs extinction of social fear

To confirm the results of the pharmacological inhibition of HDAC1, we performed a gain-of-function experiment by viral overexpression of HDAC1 in the septum. All SFC^+^ mice received the same number of CS-US pairings during acquisition of social fear (Figure 4B) three weeks after viral transfection. All SFC^+^ mice with viral overexpression of either HDAC1 or GFP exhibited high levels of social fear during the presentation of the first social stimulus, as reflected by reduced social investigation (Figure 4C). However, social fear extinction was significantly impaired by septal HDAC1 overexpression, as the SFC^+^/HDAC1 group displayed lower social investigation during exposure of the 2^nd^ to the 6^th^ social stimuls (p<0.05 vs SFC^+^/GFP; Figure 4C). In SFC^-^ mice, septal HDAC1 overexpression had no effect on social investigation times (Figure 4C-D). All treatment groups showed similarly high levels of social investigation during social fear recall on day 3 of the SFC paradigm (Figure 4D), indicating successful extinction of social fear. Overexpression of HDAC1 was confirmed by Western Blot analysis (Supplementary Figure S4B).

### Experiment 5: Pharmacological inhibition of LS-HDAC1 facilitates consolidation of long-term social fear extinction memory

In order to investigate the role of HDAC1 in enhancing the endurance of social fear extinction memory, we studied the long-term (30 days after extinction) behavioral consequences of local pre-extinction inhibition of HDAC1. During acquisition of social fear, both Veh and MS275-infused SFC^+^ mice received the same number of CS-US pairings (Figure 5B). Replicating the previous results (Figure 2), SFC^+^ mice infused with MS275 showed facilitated extinction, which can be inferred from the increased social investigation during exposure to the 2^nd^ and 3^rd^ stimuli (p<0.05 vs SFC^+^/Veh; Figure 5C). Consequently, both groups showed similar levels of social investigation times during exposure to the 6^th^ social stimulus, suggesting successful extinction of social fear. During social fear extinction recall performed 30 days after social fear acquisition, SFC^+^/MS275 mice showed significantly higher levels of social investigation (p<0.05 vs. SFC^+^/Veh; Figure 5D), which suggested that the facilitation of social fear extinction by blockade of LS-HDAC1 also results in enhanced long-lasting extinction memory formation.

## Discussion

Processes that regulate the transition from short-term to long-lasting remission in the context of SAD are not well understood. This study uses the SFC-paradigm to establish a novel epigenetic mechanisms within the LS, which specifically regulates the long-lasting extinction of socially aversive memories. Here we have identified HDAC1 as a key regulator of both acute social fear extinction and of long-lasting consolidation of social fear extinction memory within the LS of male mice. These findings are based on the observations of (i) activation of the LS in response to acquisition of social fear, (ii) increased activity-inducing phosphorylation of HDAC1 within the LS of SFC^+^ mice during the course of social fear extinction, (iii) bidirectional and specific regulation of social fear expression by either loss or gain-of-function of HDAC1 within the LS, and (iv) long-lasting consolidation of extinction memory after pharmacological inhibition of HDAC1 within the LS.

### LS is specifically activated by social fear acquisition

We found a context-specific activation of the LS after fear conditioning based on cFos expression as a reliable marker of cellular activity. The LS was found to be exclusively activated in response to acquisition of social fear (Figure 1C-D), whereas its activity remained unchanged throughout the CFC paradigm (Figure 1F-G). This result is in agreement with the recent demonstration of similar post-acquisition activation of the LS in virgin female SFC^+^ mice [22], and suggests that the LS is specifically involved in processing social information during social fear acquisition. Indeed neurotransmission within the LS has emerged as a hotspot for the regulation of stress-induced alterations in social behaviour of mice including social fear [8,22], impaired social interaction [27] and aggression [28,29]. Interestingly, the hypothalamic PVN, which is a stress-responsive brain structure, was also activated in both social (Supplementary Figure S2C) and cued (Supplementary Figure S2F) fear conditioned mice 90min after acquisition. This suggest that, although fear acquisition is in general a stressful experience, processing of socially relevant stressful information requires LS activity. This theory is supported by the fact that cFos expression in the LS is an indicator of long-term social memory in mice [30]. Hence, specific activation of LS cells upon social fear acquisition and its reduced activity during extinction could be an indication of strength of the acquisition memory. The activity patterns within the LS during the SFC and CFC paradigms were different from that of the BLA, as elevated cFos levels were observed within the BLA in response to acquisition of both social (Supplementary Figure S1B) and cued fear (Supplementary Figure S1E). Such enhanced activation of the BLA, irrespective of the nature of the stimuli, suggests that the association of CS (both social and non-social) and US requires the activation of the BLA. This is in line with previous studies suggesting the BLA to be a sensory interface for cue-specific cortical inputs to the thalamus [31,32]. Therefore, neuronal activity within the BLA seems to be required for the formation of both social and non-social aversive memories. Thus, our data in combination with pre-existing literature suggest that plasticity of the LS might play a key role in regulating the valence of socially relevant cues and in the formation of associative memories in a social context.

### Activity-inducing phosphorylation of HDAC1 is increased in the LS during the extinction of social fear

In contrast to their diverse roles in regulation of associative memory formation, preliminary analysis of septal mRNA expression revealed that of all the class I HDACs, only *Hdac1* was dynamically regulated during SFC with its peak expression observed 90min after acquisition of social fear and this increased reverted to baseline levels post social fear extinction (Supplementary Figure S2B). Based on this information, and the facts that (i) class I HDACs regulate contextual memory [14,15,17], and (ii) the LS is essentially involved in processing of social information [22,33,34], we assessed protein levels of HDAC1 and its activity-inducing phosphorylation within the LS of mice across social fear extinction. Although HDAC1 protein levels remained unchanged across extinction (Figure 2D), we found that its phosphorylation at ser421 and ser423 were enhanced within the LS of SFC^+^ across social fear extinction (Figure 2E-F). Casein kinase 2-mediated post-translational modifications of HDAC1, such as phosphorylation at ser421 and ser423, are known to regulate its catalytic (i.e., deacetylase) activity and its association with large repressor complexes, such as Sin3A and CoREST complexes [35]. Increased phosphorylation at these sites as a result of extinction of social fear, and an inverse correlation between ppHDAC1 and investigation fold change in SFC^+^ mice (Supplementary Figure S2K) suggests that extinction training increases HDAC1 activity only in fear conditioned mice. This data, along with the reduction in cFos expression after social fear extinction within the LS (Figure 1C and 1D), points towards a decrease in plasticity within the LS during social fear extinction. Indeed, HDAC1 has been previously identified as a negative regulator of cFos expression [36,37] advocating that HDAC1 might suppress neuronal activity and plasticity within the LS. In support, increased HDAC1 occupancy and consequent deacetylation of the cFos promoter result in reduced cFos expression in the hippocampus and has been correlated with impaired extinction of contextual fear [14]. Thus, we propose that HDAC1 might mediate the reduction in cFos-induced neuronal plasticity within the LS during learning of social fear extinction. Such reduction in plasticity within the LS correlating with extinction of social fear might impede not only extinction learning but also long-lasting extinction memory formation.

### HDAC1 is a negative regulator of enduring extinction of social fear in male mice

In an attempt to causally link the elevated HDAC1 activity within the LS with social fear, we found that pharmacological blockade of LS-HDAC1 120 min prior to fear extinction specifically facilitated extinction of social fear (Figure 3C). Conversely, AAV-mediated overexpression of HDAC1 within the LS impaired extinction of social fear (Figure 4C). As mentioned earlier, HDAC1 as well as other class I HDACs, have been described to be critical regulators of fear memory [14]. For example, reduced HDAC3 occupancy at the cFos promoter in the dorsal hippocampus and BLA correlated with the formation of contextual fear memory without affecting cued fear [17]. In another study, RGFP963 – a small molecule inhibitor of class I HDACs – enhanced consolidation of cued fear [19]. Considering the fact that the LS is an important brain structure for processing social cues during the formation of associative memories, our results suggest that HDAC1-mediated reduction in plasticity within the LS might specifically impede extinction of social fear.

One of the most pressing issues concerning evidence-based combinatorial therapy has been the fact that the majority of SAD patients report a relapse of aversive memories after completion of treatment. This phenomenon is also called ‘spontaneous recovery’ of fear memories in rodent models depicting anxiety disorders [18,38]. Thus, finding novel pharmacological treatment options with increased endurance of extinction memories is of utmost importance. To this end, we studied the effect of preextinction pharmacological inhibition of HDAC1 on the formation of long-lasting social fear extinction memories. We found that blockade of LS-HDAC1 was sufficient to increase the endurance of social fear extinction and prevented spontaneous recovery of aversive social memories as found in vehicle-treated controls (Figure 5D). This corroborates the idea that HDAC inhibitors are effective cognitive enhancers [18,39].

In summary, our study provides important insight into the molecular mechanisms, which are involved in regulation of social fear extinction, and implicates that HDAC1 within the LS acts as negative regulator of social fear extinction in mice. To the best of our knowledge, this is the first study to causally link HDAC1 to regulation of aversive social memories. This study adds to the growing body of evidence placing HDACs as potential regulators of associative memory and specifically propounds HDAC1 as a potential therapeutic target for SAD.

## Acknowledgements

We would like to thank Rodrigue Maloumby, Martina Fuchs, Ramona Pawlak and Verena Biermeier for their excellent technical assistance.

## Author contributions

AB and RM conceived the project, designed and performed experiments, analysed the data and prepared the manuscript. AP performed experiments and analysed data. IDN and RM provided scientific discussion, intellectual contributions and revised the manuscript.

