## Supplementary information for "Septal HDAC1 facilitates long-lasting extinction of social fear in male mice"

**Supplementary methods:**

**Social fear conditioning paradigm**

*Social fear acquisition (day 1)*. For social fear acquisition each mouse was transferred from its home cage into the conditioning chamber (45x23x36cm; transparent Perspex box with a stainless-steel grid floor). After a 30-s adaptation period, an empty wire mesh cage (7x7x6cm) was presented as a non-social stimulus for 3min, which was replaced with an identical small cage containing an unfamiliar conspecific (conditioned stimulus, CS). Non-conditioned controls (SFC**^-^**) freely investigated the social stimulus in the conditioning chamber for 3min without receiving any foot shocks, whereas SFC**^+^** mice were given a 1-s electric foot shock (0.7mA, unconditioned stimulus, US) each time they investigated the social stimulus. On average, SFC**^+^** mice received 2-3 foot shocks (mean 2.4 foot shocks/mouse) and were returned to their home cage, when no further social contact was made for 2min.

*Social fear extinction training (day 2).* One day after SFC, mice were exposed to 3 non-social stimuli (empty cages) in their home cage to assess non-social investigation as a parameter of non-social fear and general anxiety-related behavior. Mice were then exposed to 6 unfamiliar social stimuli, i.e., six different adult male mice in small cages, to assess social investigation as a parameter of social fear. Each stimulus was presented for 3min, with a 3-min inter-stimulus interval.

*Social fear extinction recall (day 3 or 31).* One day (experiment 1-4) or 30 days (experiment 5) after extinction training, mice were exposed to 6 different, unfamiliar social stimuli, each in a different small cage (see days 1 and 2), in their home cage for 3min with a 3-min inter-stimulus interval.

**Cued fear conditioning paradigm**

*Cued fear acquisition (day 1).* All mice were placed in the conditioning chamber (context A; transparent Perspex box: 23×23×36cm with electric grid floor; cleaned with lemon-scented detergent) and, after a 5-min adaptation period, exposed to five CS-US pairings with a 2-min inter-stimulus interval. The CS (80dB, 8kHz, 30s continuous sound) co-terminated with a mild electric foot shock (US; 0.7mA; pulsed current, 2s). The animals were returned to their home cage 5min after the last CS-US pairing.

*Cued fear extinction training (day 2).* On day 2, mice were placed in context B (black Perspex box: 23×23×36cm with a smooth floor; cleaned with floral smelling detergent) and, after a 5-min adaptation period, exposed to 20 CS presentations with a 5-s inter-stimulus interval. They were returned to their home cage after the last CS presentation. These CS presentations were collapsed into ten blocks with the mean freezing percentage during two CS presentations represented in each block respectively.

*Cued fear extinction retention (day 3).* One day after cued fear extinction training, animals were again placed in context B, and after a 5-min adaptation period, they were exposed to 2 CS presentations with a 2-min inter-stimulus interval. Animals were returned to their home cage after the last CS presentation. These CS presentations were then collapsed into one block with each bar representing the mean freezing percentage during 2 CS presentations.

**mRNA extraction and analysis of gene expression using quantitative RT-PCR**

Tissue micro punches were homogenized in TRI reagent (Sigma) and stored at -20°C for mRNA isolation. Total RNA was isolated using chloroform extraction followed by precipitation with isopropanol and glycogen before elution into the nuclease-free water. Quality of the isolated total RNA was assessed using nanodrop spectrophotometer. A hundred nanograms of isolated mRNA were reverse transcribed into cDNA using Super Script III first strand synthesis system (Invitrogen). Relative mRNA expression for *Hdac1* (NM_008228), *Hdac2* (NM_008229), *Hdac3* (NM_010411) *and Hdac8* (NM_027382) was measured using SYBR green (Qiagen) where ribosomal protein L13A (*Rpl13A*, NM_009438) and glyceraldehyde-3-phosphate-dehydrogenase (*Gapdh*, NM_001289726). Primer sequences used for each gene is given in supplementary table 1.

| **Gene** | **Forward Primer (5’ – 3’)** | **Reverse Primer (5’ – 3’)** |
| --- | --- | --- |
| *Gapdh* | AAGGGCTCATGACCACAG | CAGGGATGATGTTCTGGG |
| *Rpl13A* | GAGGGGCAGGTTCTGGTATTG | GGGGTTGGTATTCATCCGCT |
| *Hdac1* | ACTACGACGGGGATGTTGGA | CAGCATTGGCTTTGTGAGGG |
| *Hdac2* | AGGTGAAGGAGGTCGTAGGAA | TCTGACTTGGCTCCTTTGGG |
| *Hdac3* | ATGCCTTCAACGTGGGTGAT | CAGAAGCCAGAGGCCTCAAA |
| *Hdac8* | GCAATGAGCCCCACCGAATC | TCCACAAACCGCTTGCATCA |

**Supplementary table S1:** Primer sequence for quantitative RT-PCR

**Supplementary results:**

**SFC and CFC lead to differential activation of brain regions**

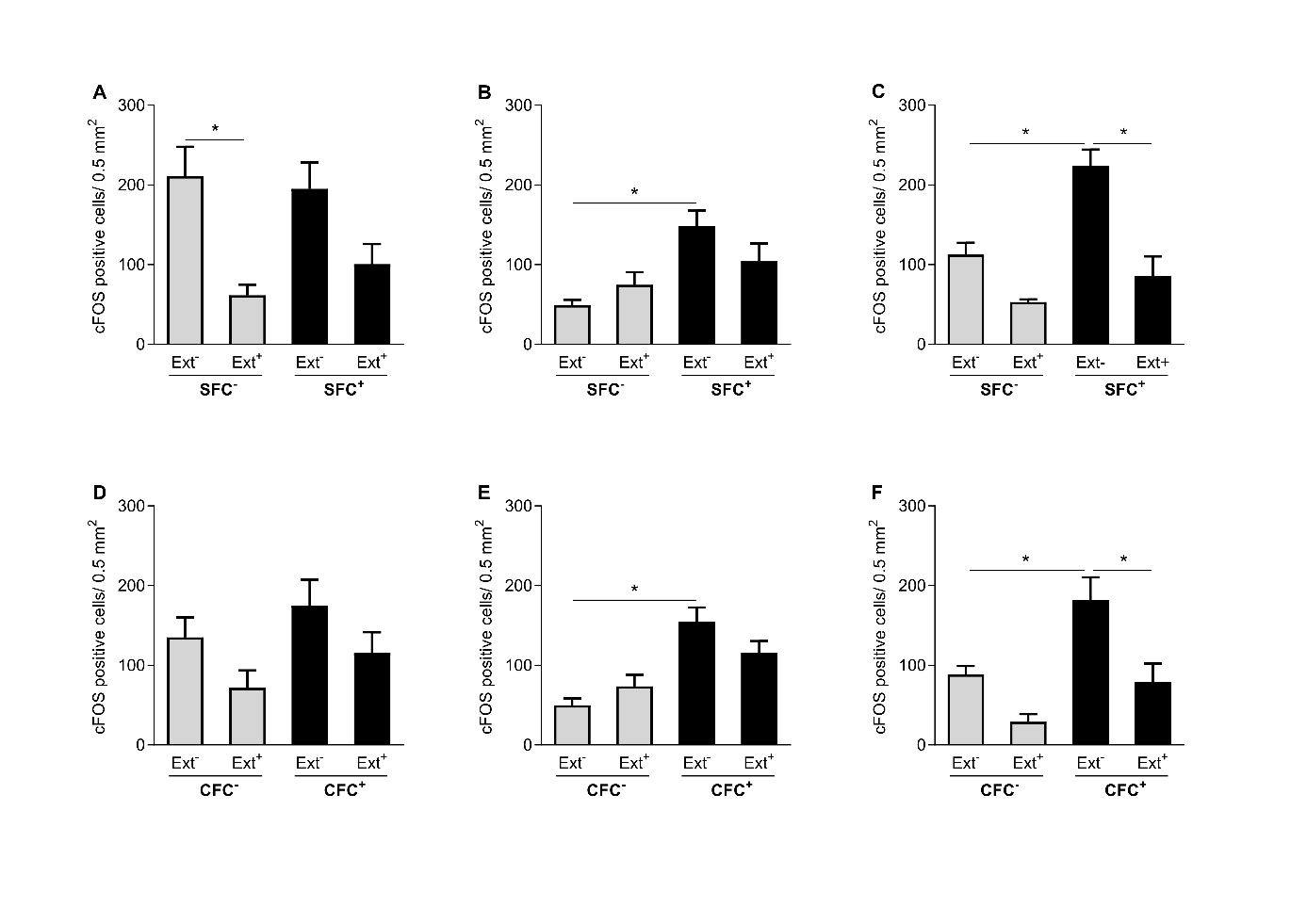

**Supplementary Figure S1:** Social fear conditioning (SFC) and cued fear conditioning (CFC) leads differential activation of ventral lateral septum (vLS; A and D), basolateral amygdala (BLA; B and E), paraventricular nucleus (PVN; C and F). Number of cFos-immunopositive cells were counted after SFC (in vLS: A, BLA: B and PVN: C) and CFC (in vLS: D, BLA: E and PVN: F). Mice were sacrificed 90 min after, either social fear acquisition (SFC**^-^** / Ext**^-^** and SFC**^+^** / Ext**^-^**; n = 4-6) and after cued fear acquisition (CFC**^-^** / Ext**^-^** and CFC**^+^**/ Ext**^-^**; n = 4-6) on day 1 or after social fear extinction (A; SFC**^+^** / Ext**^+^** and SFC**^+^** / Ext**^+^**; n = 4-6) and after cued fear acquisition (D; CFC**^-^** / Ext**^+^** and CFC**^+^** / Ext**^+^**; n = 4-6) on day 2. Data represents mean number of cFos immunopositive cells/ 0.5 mm^2^ ± SEM. *p<0.05 vs SFC**^-^**  between indicated groups.

**SFC leads to differential expression of HDAC within the LS and ppHDAC1 levels within the LS correlates with investigation fold change across extinction**

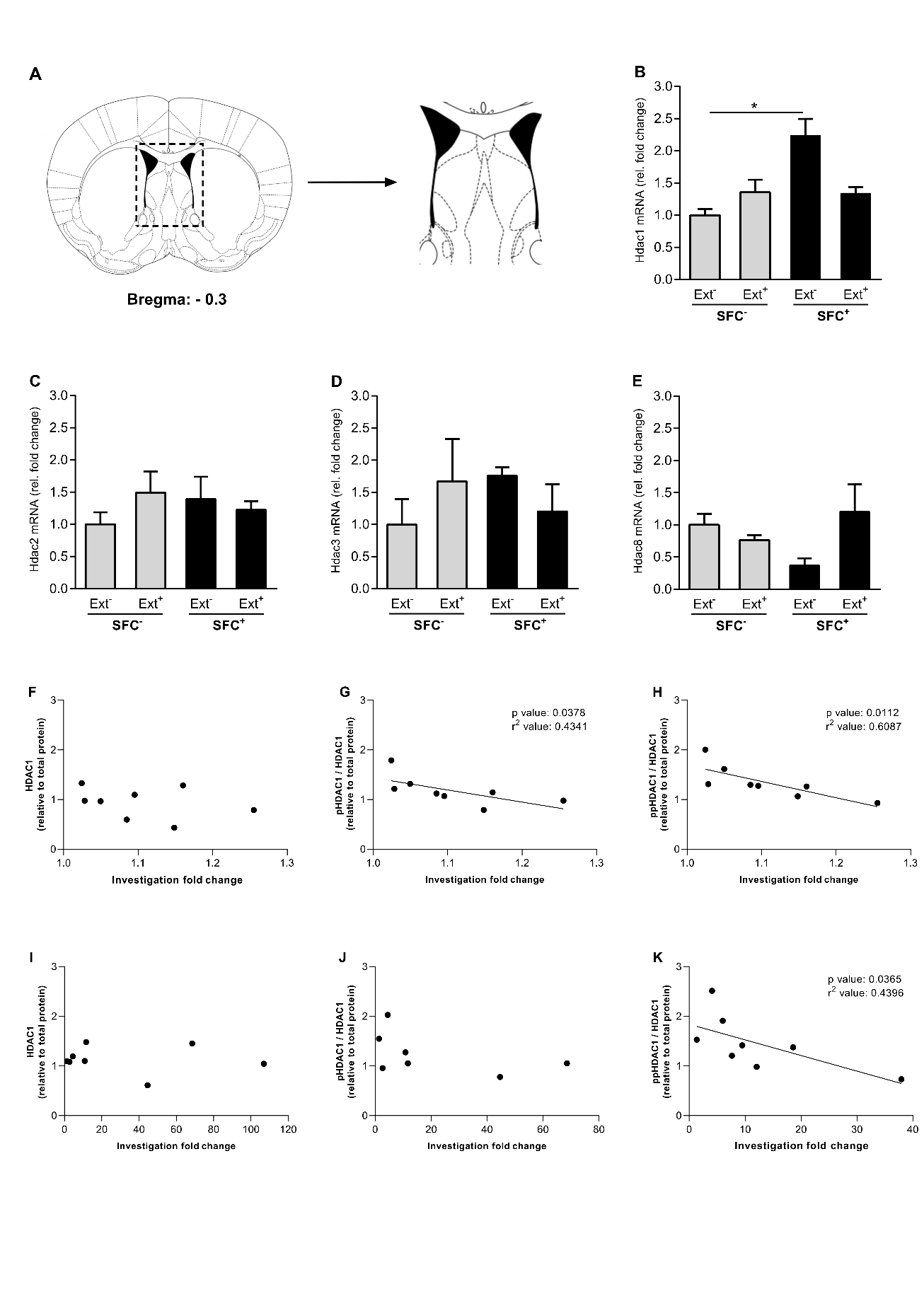

**Supplementary Figure S2:** Hdac1 mRNA is specifically upregulated after social fear conditioning within the mice LS (B-E). Schematic representation of the are from which the tissue micropunches were obtained (A). mRNA was isolated from mice LS 90 min after social fear conditioning (SFC**^-^**/Ext**^-^** and SFC**^+^**/Ext**^-^**; n = 6; Day 1) or after extinction (SFC**^-^**/Ext**^+^** and SFC**^+^**/Ext**^+^**; n = 6; Day 2) and gene expression for Hdac1 (B), Hdac2 (C), Hdac3 (D), and Hdac8 (E) was measured. In a separate set of mice, level of HDAC1 and its phosphorylations (i.e. ser421: pHDAC1, ser421 and ser423: ppHDAC1) within the septum measured by Western blot was correlated with investigation fold change across social fear extinction. Mice were social fear conditioned on day 1 of the SFC paradigm. Total protein was isolated from mice septum 90 min after extinction and correlations were performed between investigation fold change for SFC**^-^** mice and level of HDAC1 (F), pHDAC1 (G) and ppHDAC1 (H), or investigation fold change for SFC^+^ mice and level of HDAC1 (I), pHDAC1 (J) and ppHDAC1 (K). Data represent mean fold change + SEM (B-E), correlation between investigation fold change and individual protein levels (F-K).

**Pre-extinction pharmacological blockade of HDAC1 within the LS does not effect extinction of cued fear**

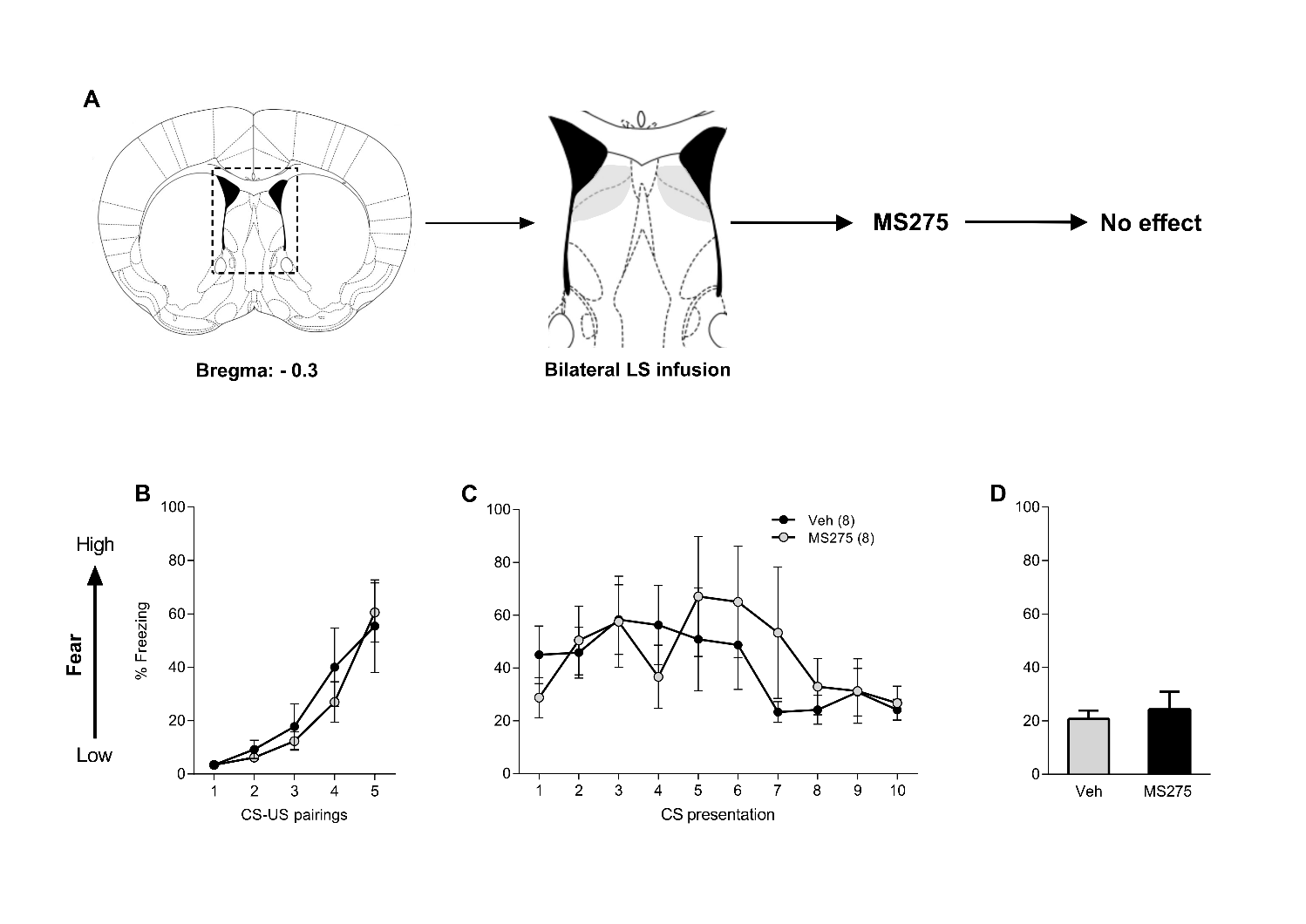

**Supplementary Figure S3:** Pharmacological blockade of HDAC1 within the lateral septum (LS) does not effect cued fear. Schematic representation of infusions (A). Male mice were cued fear conditioned (Day 1; B) followed by cued fear extinction (Day 2; B) and cued fear extinction recall (Day 3; C) during which percentage freezing was measured. Fear conditioned mice were bilaterally infused within the LS with Veh (Ringer / 0.2µl / side) or MS275 (375ng/ 0.2µl/ hemisphere) into the LS 60 min prior to extinction training. Data represent mean percentage of freezing ± SEM (B-D).

**AAV mediated genetic manipulation leads to overexpression of HDAC1**

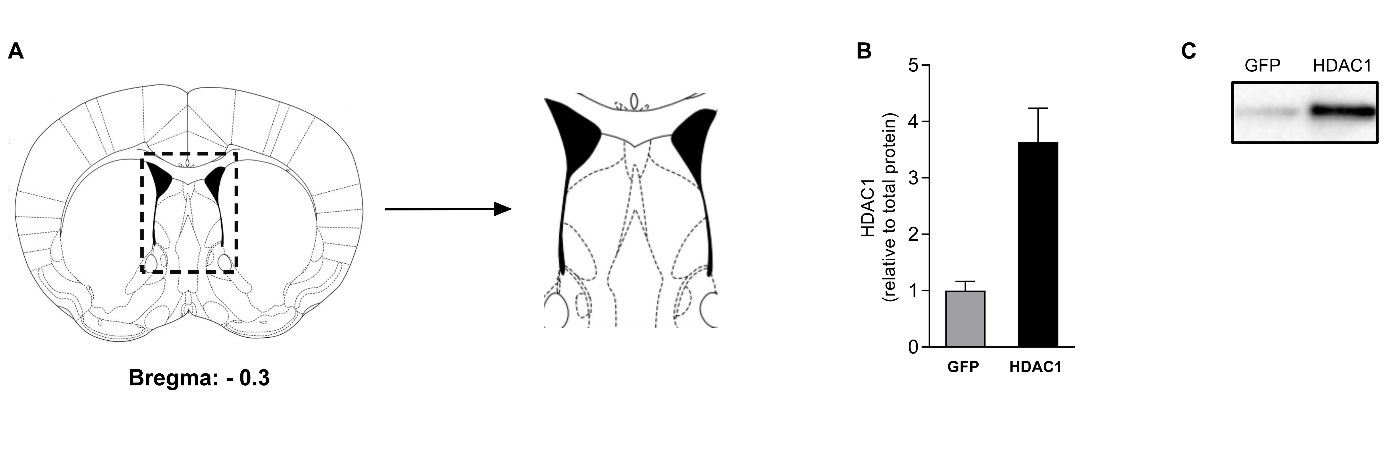

**Supplementary Figure S4:** Genetic manipulation leads to overexpression of HDAC 1. Schematic representation of the septum including viral manipulations (A). The highlighted area was considered as adequate infusion localization. Male mice were injected with AAV-hSyn-HDAC-GFP-WPRE (HDAC1) or AAV-hSyn-GFP-WPRE (GFP; control) 3 weeks prior to acquisition of social fear. After the behavioral analysis using the SFC paradigm, HDAC1 protein levels were measured from septum micropunches using Western blot (B). Representative western blot images for HDAC1 (C). Data represent mean fold change + SEM (D-F). *p<0.05 HDAC1 *vs* GFP (B).

**Statistics table:**

| **Experiment 1. The LS is specifically activated by SFC, but not CFC (Figure 1).** | | |
| --- | --- | --- |
| *SFC*  Social fear extinction (B)  *cFos immunohistochemistry*  cFos positive cells in dLS (C)  *CFC*  Cued fear extinction (E)  *cFos immunohistochemistry*  cFos positive cells in dLS (F) | *Group effect (SFC)*  F(1, 10) = 11.25, p = 0.0073*  *Group effect (SFC x extinction)*  F(3, 15) = 13.10, p = 0.0002*  *Group effect (CFC)*  F(1, 100) = 172.4, p < 0.0001*  *Group effect (SFC x extinction)*  F(3, 17) = 0.3006, p = 0.0.8245 | *Group x extinction effect*  F(8, 80) = 7.222, p < 0.0001*  *Group x extinction effect*  F(9, 100) = 22.78, p < 0.0001* |
| **Experiment 2: HDAC1 phosphorylation is upregulated in response to social fear extinction within the septum of fear conditioned mice (Figure 2).** | | |
| *SFC*  Social fear acquisition (A)  Social fear extinction: S1 (B)  Social fear extinction: S1-6 (C)  *Western Blot*  HDAC1 (D)  pHDAC1 (E)  ppHDAC1 (F) | *Group effect (SFC)*  T(17) = 0.9732, p = 0.4015  F(1, 17) = 26.99, p < 0.0001*  F(1, 16) = 43.96, p < 0.0001*  *Group effect (SFC x extinction)*  F(3, 31) = 0.7825, p = 0.5128  F(3, 33) = 3,842, p = 0.0183*  F(3, 32) = 4.114, p = 0.0141* | *Group x extinction effect*  F(3, 51) = 41.90, p < 0.0001*  F(8, 128) = 11.47, p < 0.0001* |
| **Experiment 3: Pharmacological inhibition of LS-HDAC1 facilitates extinction of social fear (Figure 3).** | | |
| *SFC (MS275 vs Veh)*  Social fear acquisition (B)  Social fear extinction (C)  Social fear recall (D) | *Group effect (Treatment x SFC)*  T(16) = 0.9177, p = 0.3724  F(3, 234) = 17.30, p < 0.0001*  F(3, 156) = 5.268, p = 0.0017* | *Group x extinction effect*  F(24, 234) = 2.619, p = 0.0001*  F(15, 156) = 0.1601, p = 0.9999 |
| **Experiment 4: AAV-mediated overexpression of HDAC1 within the LS impairs extinction of social fear (Figure 4).** | | |
| *SFC (HDAC1 vs GFP)*  Social fear acquisition (B)  Social fear extinction (C)  Social fear recall (D) | *Group effect (Treatment x SFC)*  T(18) = 0.8847, p = 0. 3880  F(3, 30) = 45.40, p < 0.0001*  F(3, 31) = 2.941, p = 0.0484* | *Group x extinction effect*  F(24, 240) = 8.131, p < 0.0001*  F(15, 155) = 1.129, p = 0.3348 |
| **Experiment 5: Pharmacological inhibition of LS-HDAC1 facilitates consolidation of long-term social fear extinction memory (Figure 5).** | | |
| *SFC*  Social fear acquisition (B)  Social fear extinction (C)  Social fear recall (D) | *Group effect (SFC)*  T(16) = 0.6976, p = 0.4954  F(1, 16) = 6.892, p = 0.0184*  T(17) = 2.534, p = 0.0214* | *Group x extinction effect*  F(8, 128) = 1.767, p = 0.0895 |
| **Figure S1: SFC and CFC leads differential activation of vLS, BLA and PVN.** | | |
| *cFos immunohistochemistry*  SFC  cFos positive cells in vLS (A)  cFos positive cells in BLA (B)  cFos positive cells in PVN (C)  CFC  cFos positive cells in vLS (D)  cFos positive cells in BLA (E)  cFos positive cells in PVN (F) | *Group effect (SFC)*  F(3, 16) = 5.633, p = 0.0079*  F(3, 13) = 6.275, p = 0.0072*  F(3, 11) = 17.36, p = 0.0002*  F(3, 17) = 2.535, p = 0.0912  F(1, 18) = 11.40, p = 0.0002*  F(2, 13) = 6.271, p = 0.0124* |  |
| **Figure S2: SFC leads to differential expression of HDAC within the LS and ppHDAC1 levels within the LS correlates with investigation fold change across extinction.** | | |
| *qRT-PCR*  Hdac1 mRNA (B)  Hdac2 mRNA (C)  Hdac3 mRNA (D)  Hdac8 mRNA (E)  *Investigation fold change – Protein expression correlation*  SFC^-^  - HDAC1 (F)  SFC^-^  - pHDAC1 (G)  SFC^-^  - ppHDAC1 (H)  SFC^+^  - HDAC1 (F)  SFC^+^  - pHDAC1 (G)  SFC^+^  - ppHDAC1 (H) | *Group effect (SFC x extinction)*  F(3, 39) = 9.5970, p < 0.0001*  F(3, 13) = 0.6942, p = 0.5718  F(3, 13) = 0.6668, p = 0.5873  F(3, 14) = 2.082, p = 0.2487  R^2^ = 0.1038, p = 0.2183  R^2^ = 0.4341, p = 0.0378*  R^2^ = 0.6078, p = 0.0112*  R^2^ = 0.0009, p = 0.4116  R^2^ = 0.2397, p = 0.1324  R^2^ = 0.4396, p = 0.0365* |  |
| **Figure S3: Pre-extinction pharmacological blockade of HDAC1 within the LS does not effect extinction of cued fear.** | | |
| *CFC (MS275 vs Veh)*  CFC – acquisition (B)  CFC – extinction (C)  CFC – extinction recall (D) | *Group effect (CFC)*  F(1, 14) = 0.1264, p = 0.7274  F(1, 14) = 0.0709, p = 0.7938  T(14) = 0.4648, p = 0.6492 | *Group x stimulus effect*  F(4, 56) = 0.4909, p = 0.7424  F(9, 126) = 1.366, p = 0.2106 |
| **Figure S4: AAV mediated genetic manipulation leads to overexpression of HDAC1.** | | |
| *Western blot*  HDAC1 overexpression (B) | *Group effect (GFP vs HDAC1)*  T(32) = 4.243, p = 0.0002* |  |
